# Identification and quantification of Lyme pathogen strains by deep sequencing of outer surface protein C (*ospC*) amplicons

**DOI:** 10.1101/332072

**Authors:** Lia Di, Zhenmao Wan, Saymon Akther, Chunxiao Ying, Amanda Larracuente, Li Li, Chong Di, Roy Nunez, D. Moses Cucura, Noel L. Goddard, Konstantino Krampis, Wei-gang Qiu

## Abstract

Mixed infection of a single tick or host by Lyme disease spirochetes is common and a unique challenge for diagnosis, treatment, and surveillance of Lyme disease. Here we describe a novel protocol for differentiating Lyme strains based on deep sequencing of the hypervariable outer-surface protein C locus (*ospC*). Improving upon the traditional DNA-DNA hybridization method, the next-generation sequencing-based protocol is high-throughput, quantitative, and able to detect new pathogen strains. We applied the method to over one hundred infected *Ixodes scapularis* ticks collected from New York State, USA in 2015 and 2016. Analysis of strain distributions within individual ticks suggests an overabundance of multiple infections by five or more strains, inhibitory interactions among co-infecting strains, and presence of a new strain closely related to *Borreliella bissettiae*. A supporting bioinformatics pipeline has been developed. With the newly designed pair of universal *ospC* primers targeting intergenic sequences conserved among all known Lyme pathogens, the protocol could be used for culture-free identification and quantification of Lyme pathogens in wildlife and clinical specimens across the globe.

## Introduction

Lyme disease occurs throughout the Northern Hemisphere and is the most prevalent vector-borne diseases in the United States (Hengge et al., 2003; Margos et al., 2011; Schwartz et al., 2017). The causative agents of Lyme disease are obligate bacterial parasites of vertebrates transmitted predominantly by hard-bodied *Ixodes* ticks. Lyme pathogens and related strains, formerly known as the *Borrelia burgdorferi sensu lato* species group, have been recently (and controversially) classified as a new spirochetal genus *Borreliella* (Adeolu and Gupta, 2014; Margos et al., 2017). In the US, while *B. burgdorferi* causes the majority of the Lyme disease cases, more than a half dozen additional *Borreliella* species have been recognized including *B. americana, B. andersonii, B. bissettiae, B. californiensis, B. carolinensis, B, kurtenbachii*, and *B. mayonii* (Dolan et al., 2016; Marconi et al., 1995; Margos et al., 2016, 2013; Bobbi S. Pritt et al., 2016; Rudenko et al., 2011, 2009). *Borreliella* species vary not only in genomic sequences but also in geographic distribution, host preferences, human pathogenicity, and disease manifestations (Barbour et al., 2009; Casjens et al., 2018; Kurtenbach et al., 2006; Margos et al., 2011; Mongodin et al., 2013; Bobbi S Pritt et al., 2016). In addition, the *Ixodes* ticks in the US and elsewhere are frequently co-infected with *Borrelia miyamotoii*, a member of the re-defined *Borrelia* genus now consisting exclusively of strains grouped with agents of relapsing fever (Barbour et al., 2009; Wagemakers et al., 2015).

A hallmark of Lyme disease endemics is the coexistence of multiple spirochete species and strains within local populations and oftentimes within a single vector, host or patient (Brisson and Dykhuizen, 2004; Durand et al., 2017; Guttman et al., 1996; Qiu, 2008; Seinost et al., 1999; Walter et al., 2016; Wang et al., 1999; Wormser et al., 2008). High genetic diversity within local pathogen populations is to a large extent driven and maintained by frequency-dependent selection under which rare strains gain selective advantage over common ones in establishing super-infection in a host (Bhatia et al., 2018; Durand et al., 2017; Haven et al., 2011; States et al., 2014). In addition, local con-specific strains may have diverged in host specificity and other phenotypes including human virulence and invasiveness (Brisson and Dykhuizen, 2004; Hanincova et al., 2013; Seinost et al., 1999; Wormser et al., 2008). Against this backdrop of the vast geographic, genetic, and phenotypic variations of Lyme disease pathogens across the globe and within endemic regions, it is essential to develop accurate, sensitive, and scalable technologies for identifying species and strains of Lyme pathogens in order to understand, monitor, and control the range expansion of Lyme disease (Kilpatrick et al., 2017; Kurtenbach et al., 2006; Qiu and Martin, 2014).

Early molecular technologies for identifying Lyme pathogen strains relied on amplifying and detecting genetic variations at single variable locus including the outer-surface protein A locus (*ospA*), outer surface protein C locus (*ospC*), and the intergenic spacer regions of ribosomal RNA genes (*rrs-rrlA* and *rrfA-rrlB*) (Guttman et al., 1996; Wang et al., 2014, 1999). Availability of the first Lyme pathogen genome facilitated development of more sensitive multilocus sequence typing (MLST) technologies targeting genetic variations at a set of single-copy housekeeping genes (Fraser et al., 1997; Hanincova et al., 2013; Qiu, 2008). For direct identification of Lyme strains in tick and host specimen without first culturing and isolating the organisms, a reverse-line blotting (RLB) technology has been developed based on DNA-DNA hybridization (Brisson and Dykhuizen, 2004; Durand et al., 2015; Morán Cadenas et al., 2007; Qiu et al., 2002). The RLB technology, while sensitive and able to detect mixed infection in tick and hosts, is difficult to scale up or to standardize and does not yield quantitative measures of strain diversity. A further limitation of the RLB technology is that it depends on oligonucleotide probes of known *ospC* major-group alleles and is not able to detect strains with novel *ospC* alleles.

Next-generation sequencing (NGS) technologies circumvent the limitations of traditional methods in scalability, standardization, and ability for *de novo* strain detection while offering high sensitivity and high throughput quantification (Lefterova et al., 2015). Using the hybridization capture technology to first enrich pathogen genomes in ticks and subsequently obtaining genome-wide short-read sequences using the Illumina NGS platform, >70% of field-collected nymphal ticks from Northeast and Midwest US are found to be infected with multiple *B. burgdorferi* strains due to mixed inoculum (Walter et al., 2016). In an NGS-based study of European Lyme pathogen populations, a combination of quantitative PCR and high-throughput sequencing on the 454 pyrosequencing platform targeting the *ospC* locus and revealed a similarly high rate (77.1%) of mixed infection of nymphal ticks by *B. afzelii* and *B. garinii* (Durand et al., 2017).

Here we report an improved NGS technology for identifying Lyme pathogen strains through deep sequencing of *ospC* sequences amplified from individual ticks. We applied the technology to over 100 pathogen-infected *Ixodes scapularis* ticks collected from New York State during a period of two years. Our results suggest a new putative *Borreliella* species, competitive interactions among co-infecting strains, and genetic homogeneity within an endemic region.

## Materials & Methods

### Tick collection and DNA extraction

Adult and nymphal blacklegged ticks (*Ixodes scapularis*) were collected in 2015 and 2016 during their host-questing seasons from four locations in endemic areas of Lyme disease surrounding New York City (Figure 1). Ticks are stored at −80°C before dissection. Each tick is immersed in 5% solution of Chelex 100 resin (Sigma-Aldrich, St. Louis, MO, USA) containing 20mg/ml Proteinase K in milliQ water (EMD Millipore, Billerica, MA, USA) with a total volume of 30μl for nymphs, 100μl for males, and 200μl for females. Ticks are dissected into four or more pieces using sterilized scalpel or disposable pipette tips. The dissected mixture is incubated at 56°C overnight and heated to 100°C for 10 minutes afterwards in a dry bath, and then briefly centrifuged to separate the tick debris and Chelex resin from the supernatant. The supernatant containing the extracted DNA is transferred to a fresh tube and stored at 4°C (or frozen at −20°C for long term storage).

**Figure 1.**
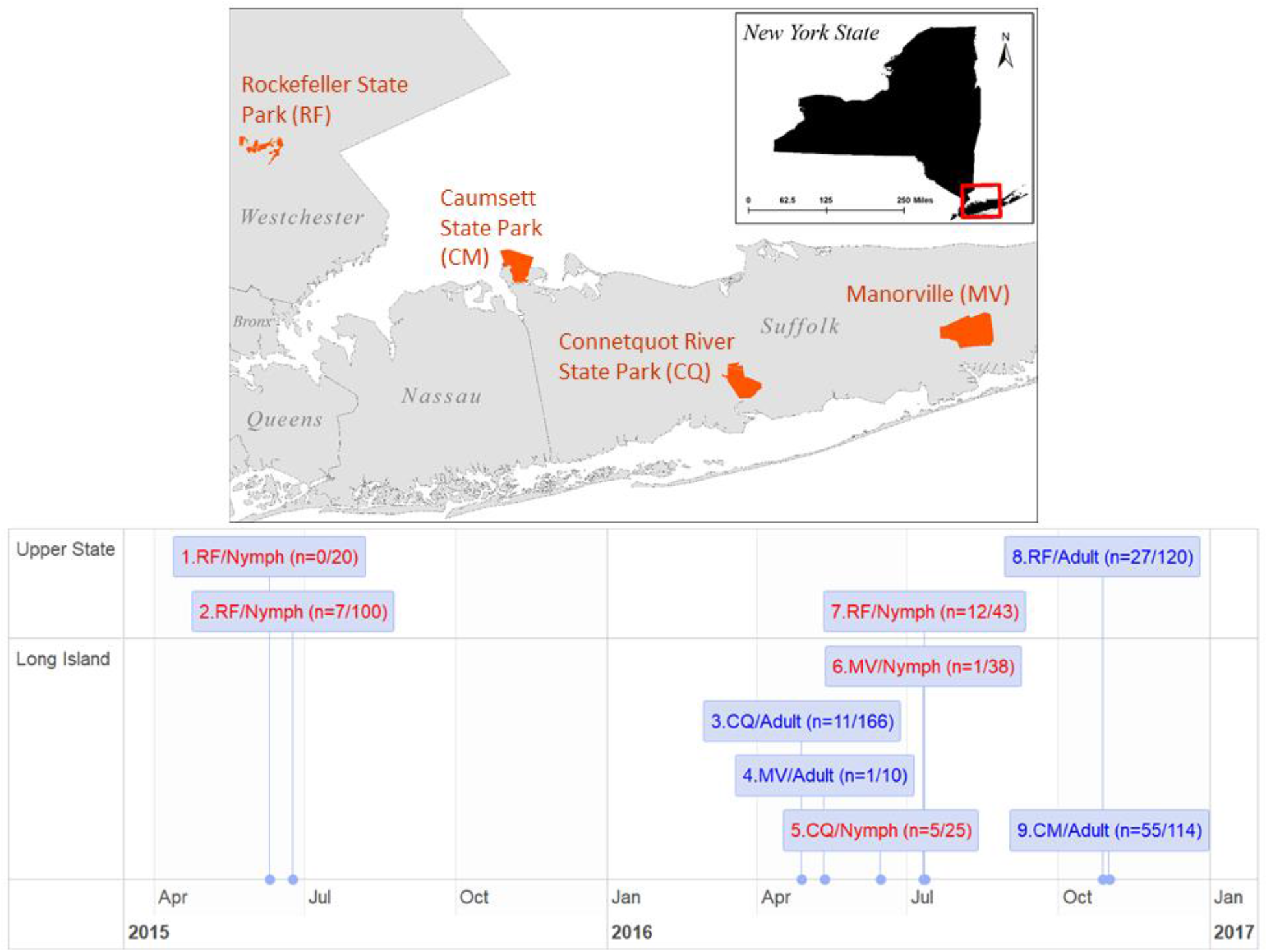
Study sites and timeline. Adult and nymphal *Ixodes scapularis* ticks were collected from four study sites in New York State, US (*top*) during their host-questing seasons in a period of 18 months (*bottom*). Nymphal samples are colored in red and adult samples in blue. Numbers in parenthesis indicate the number of ticks infected by Lyme disease spirochetes (numerator) and the total sample size (denominator).

### Single-round PCR amplification of full-length *ospC*

An improved protocol for amplifying *ospC* sequences from tick DNA extracts has been developed. First, this protocol is simpler with a single instead of two rounds of polymerase-chain reaction (Brisson and Dykhuizen, 2004; Qiu et al., 2002). Second, using a newly designed oligonucleotide primer pair targeting flanking intergenic sequences conserved across *Borreliella* species, we are able to amplify full-length (~718 bp) *ospC* sequences from all strains. Third, the new primers are able to amplify a *vsp* locus in the *B. miyamotoii* genome, enabling co-detection of *Borreliella* species and *Borrelia miyamotoii*, two major groups of Lyme pathogens in Northeast US (Barbour et al., 2009). The new primer sequences are 5’-AATAAAAAGGAGGCACAAATTAATG-3’ (“Oc-Fwd”, targeting the intergenic spacer between BB_B18 and BB_B19) and 5’-ATATTGACTTTATTTTTCCAGTTAC-3’ (“Oc-Rev”, targeting the intergenic spacer between BB_B19 and BB_B22). Alignments of primer regions for Lyme pathogens are provided as Supplemental Material S1.

Each 20μl reaction mixture contains 200 μM of each dNTP, 1U Roche FastStart Taq DNA polymerase (Roche Diagnostics, Mannheim, Germany), 2μl of 10x Roche FastStart Buffer (Roche Diagnostics, California, USA), 0.4μM of each primer and 1μl DNA extract. The reaction mixture is heated at 95°C for 4 minutes, then amplified for 36 cycles at 95°C for 30 seconds, 58°C for 30 seconds, and 72°C for 60 seconds, and finally incubated at 72°C for 5 minutes. The PCR products are electrophoresed on a 1% agarose gel, stained with ethidium bromide, and imaged under a UV light. Agencourt AMPure XP PCR Purification Kit (Beckman Coulter, Brea, CA, USA) is used to remove excess primers, dNTPs, and other reagents. Amplican quantity is measured on the Qubit 4 Fluoreometer (Thermo Fisher Scientific, Waltham, MA, USA) using the companying Qubit dsDNA HS Assay Kit.

### NGS library preparation and short-read sequencing

We followed the Nextera XT DNA Library Prep (Illumina, CA, USA, catalog no. FC-131-1024) protocol to prepare the amplicon libraries for sequencing. First, we dilute the PCR products to 0.2ng/μl after DNA quantification using a DNA 1000 kit on a 2100 BioAnalyzer (Agilent, Santa Clara, CA, USA). Samples are tagmented by incubation of 5μl DNA sample in 55°C for 5 minutes in a solution containing 10μl Tagment DNA Buffer and 5μl Amplicon Tagment Mix. Tagmentation reaction is terminated by adding 5μl Neutralize Tagment Buffer. Tagmented samples are amplified and barcoded (with Set A and Set B) using PCR in a solution containing 5μl of each barcoded primers and the Nextera PCR Mater Mix. The thermal cycling parameters are incubation at 72°C for 3 minutes, 95°C for 30 seconds, 12 cycles of 95°C for 10 seconds, 55°C for 30 seconds, and 72°C for 30 seconds, and a final incubation at 72°C for 5 minutes. The indexed amplicon libraries are cleaned using AMPure XP PCR Purification Kit and concentrations quantified using the High-Sensitivity DNA 1000 Kit on a 2100 BioAnalyzer. Amplicon libraries are diluted to the same concentration and then combined to a total concentration of 2 nM to 4 nM with a volume of 5μl or more.

In preparation for loading on the MiSeq sequencer, the pooled library is denatured by mixing 5μl of 0.2N NaOH with 5μl of sample and incubating at room temperature for 5 minutes and then adding 990μl pre-chilled Hybridization Buffer, resulting in a total of 1 ml (10pM concentration) of denatured pooled amplicon library (Illumina Denature and Dilute Libraries Guide pub. no. 15039740). Furthermore, 5% PhiX Sequencing Control (Illumina, CA, USA, catalog no. FC-110-3001) is added to the samples pool before loading to the MiSeq. The sequencing kit used is the MiSeq Reagent Kit v3 for 150 cycles (Illumina, CA, USA, catalog no. MS-102-3001), for paired reads of 75 bases each. Following sequencing, a total of 4.24 gigabases of sequence are generated by the instrument, corresponding to 57,463,220 reads, with approximately 90% of the reads (52,311,968) passing the filter build-in the MiSeq for quality control. Finally, the samples are automatically de-multiplexed to individual FASTQ files following completion of the sequencing run by the MiSeq Reporter Software based on the Nextera XT barcodes corresponding to each sample (the barcodes are also trimmed by the software from each read).

### Amplicon cloning & Sanger sequencing

New alleles are identified when the majority of reads are not aligned to reference sequences. For such samples, we performed *de novo* assembly of short reads to obtain candidate allele sequences (see below). The novel alleles are subsequently validated by cloning and Sanger sequencing. Cloning of PCR products is performed using the TOPO TA Cloning Kit for Sequencing (Thermo Fisher Scientific, Waltham, MA, USA) following manufacturer’s protocol. Five bacterial colonies containing plasmids with PCR amplicon as inserts are selected for further growth in selective liquid media. Plasmid DNA is extracted and purified using the PureLink Quick Plasmid Miniprep Kits (Thermo Fisher Scientific, Waltham, MA, USA). Nucleotide sequences of cloned PCR amplicons are obtained using the Sanger method through commercial sequencing services including Genewiz (South Plainfield, NJ, USA) and Macrogen (Rockville, MD, USA).

### Bioinformatics methods for allele identification and quantification

Alleles present in tick samples are identified and quantified by aligning the paired-end short reads to a set of reference sequences. These reference sequences consist of full-length *ospC* sequences and are obtained from published genome sequences, from Sanger sequencing of cloned or uncloned amplicons (see above), or from *de novo* assembly of short reads (see below) (the 20 reference sequences are listed in Supplemental Material S2).

The short reads are indexed and aligned to the reference sequences using software packages bwa (Li and Durbin, 2009) and samtools (Li et al., 2009). Coverage of reads at each site of each reference sequence is obtained by using bedtools (Quinlan and Hall, 2010) and visualized using ggplot2 in the R statistical computing environment (R Core Team, 2013; Wickham, 2009). Presence of new alleles is noted when a large number of reads are unmapped. For these samples, we cloned the PCR amplicons and sequenced the clones using Sanger sequencing (see above). A number of know alleles do not have full-length *ospC* sequences from sequenced genomes or from GenBank. For these alleles, we performed *de novo* assembly of reads to obtain the 5’- and 3’-end sequences using the assembler metaSPAdes (Nurk et al., 2017).

To test the accuracy and sensitivity of our bioinformatics pipelines, we generated simulated short reads with known allele identities and known proportions of allele mixture using wgsim, a part of the software samtools package (Li et al., 2009). Key steps and commands for allele identification, coverage calculation, *de novo* assembly, and simulated reads are provided as Supplemental Material S3.

### Statistical analysis of genetic diversity

We estimate the relative amount of spirochete load in individual infected ticks by the weight (in ng) of PCR amplicons. Diversity of *ospC* alleles in individual infected ticks is first measured with multiplicity, i.e., the number of unique alleles present in a sample. Allelic diversity is further measured with the Shannon diversity index 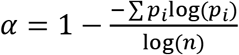, where *p_i_* is the frequency of allele *i* and *n* is the number of distinct alleles in an infected tick (allowing α=0 for *n*=1). This Shannon diversity index, also known as the Shannon Equitability Index, is a normalized measure of biodiversity ranging from 0 (infected with a single strain) to 1 (all strains being equally frequent) (Vidakovic, 2011). Allele frequencies in an infected tick are obtained as 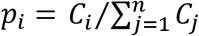, where *C_i_* is the coverage of allele *i* averaged over all nucleotide positions.

Genetic differentiation between two populations (A and B) is measured with the *F_st_* statistics: 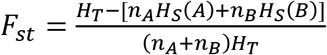 (Nei, 1973), where *H_S_*(*A*), *H_S_*(*B*), and *H_T_* are heterozygosity of sample A, sample B, and the total sample, respectively, and *n_A_* and *n_B_* are the sample sizes. Statistical significance of an *F_st_* value is estimated by a randomization procedure by which the two population samples are combined and randomly divided into two pseudo-samples with the same sample sizes. An *F_st_* value is calculated between the two pseudo-samples. The procedure is repeated for 999 times and a *p*-value is obtain as the proportion of permutated *F_st_* values that is greater than or equal to the observed value. Genetic differentiation is further tested using F-statistics implemented in the *hierfstat* package on the R statistical computing environment (Goudet, 2005).

### Data availability

New sequences have been deposited in GenBank with consecutive accessions MH071430 through MH071436. Experimental data are stored in a custom relational database. An interactive website has been developed using the D3js (http://d3js.org) JavaScript library to visualize allele composition and read depth for the 119 tick samples and is publicly available at http://diverge.hunter.cuny.edu/~weigang/ospC-sequencing/. Data sets and R scripts are publicly available at Github https://github.com/weigangq/ocseq.

## Results

### Tick infection rates, co-infections, specificity, and sensitivity

Approximately 25% of nymphal ticks and 50% of adult ticks are infected with *Borreliella* species or *Borrelia miyamotoi*. For example, the nymphal infection rate for *Borreliella burgdorferi* is 27.9% (with a 95% confidence interval of 15.3 – 43.7%) in Sample #7 and the adult infection rate for *Borreliella burgdorferi* is 42.1% (32.9 – 51.7%) for Sample #9 (Figure 1). The infection rate for *Borrelia miyamotoi* in adult ticks is 6.1% (7 out of 114 ticks; 2.5 – 12.2% confidence interval) for Sample #9. Four ticks in Sample #9 are infected with both *Borreliella burgdorferi* and *Borrelia miyamotoi* (co-infection rate 3.5%, 0.96-8.74%). Rates from other samples are underestimates due to lack of success in tick storage, processing, DNA amplification, and NGS sequencing during protocol development. These rates are consistent with results from other studies conducted in the same region and appear to be stable through recent decades (Qiu et al., 2002; States et al., 2014).

The number of sequencing averages ~108,000 reads per tick sample. The coverage (i.e., read depth) of an allele depends on the total number of tick samples in a pooled library and the number of alleles present in a tick. Alleles are identified if the reads cover all nucleotide positions of a reference allele and the total read percentage is at least 1% of the most abundant alleles. The total sample of 119 successfully sequenced ticks are divided into four sub-population samples according to geographic origin and life stage, with allele counts of pathogens in each of the four populations listed in Table 1.

**Table 1.** Allele counts^a^

| A | B | B3 | C | C14 | D | E | F | G | H | I | J | K | L | M | N | O | T | U | vsp | total |
| --- | --- | --- | --- | --- | --- | --- | --- | --- | --- | --- | --- | --- | --- | --- | --- | --- | --- | --- | --- | --- |
| From Upper State, N=27 infected adult ticks (Pop 1) |  |  |  |  |  |  |  |  |  |  |  |  |  |  |  |  |  |  |  |  |
| 2 | 4 | 0 | 7 | 0 | 1 | 6 | 0 | 7 | 2 | 3 | 0 | 3 | 4 | 2 | 5 | 0 | 5 | 4 | 2 | 57 |
| From Upper State, N=19 infected nymphs (Pop 2) |  |  |  |  |  |  |  |  |  |  |  |  |  |  |  |  |  |  |  |  |
| 3 | 3 | 0 | 1 | 1 | 1 | 4 | 1 | 1 | 3 | 0 | 0 | 2 | 1 | 2 | 2 | 2 | 2 | 0 | 3 | 32 |
| From Long Island, N=67 infected adult ticks (Pop 3) |  |  |  |  |  |  |  |  |  |  |  |  |  |  |  |  |  |  |  |  |
| 12 | 8 | 1 | 4 | 0 | 5 | 12 | 18 | 16 | 10 | 14 | 4 | 17 | 0 | 8 | 8 | 8 | 24 | 13 | 10 | 192 |
| From Long Island, N=6 infected nymphs (Pop 4) |  |  |  |  |  |  |  |  |  |  |  |  |  |  |  |  |  |  |  |  |
| 3 | 0 | 0 | 1 | 0 | 0 | 3 | 3 | 2 | 2 | 2 | 3 | 1 | 0 | 3 | 3 | 3 | 0 | 1 | 4 | 34 |
| Sums |  |  |  |  |  |  |  |  |  |  |  |  |  |  |  |  |  |  |  |  |
| 20 | 15 | 1 | 13 | 1 | 7 | 25 | 22 | 26 | 17 | 19 | 7 | 23 | 5 | 15 | 18 | 13 | 31 | 18 | 19 | 315 |

<sup>a</sup>See Figure 5 for allele frequency distributions

Specificity of allele identification is tested by generating simulated reads from a single reference sequence and aligning the simulated reads to all reference sequences. This simulation-based test shows that the bioinformatics protocol for allele identification is highly specific, with only a small fraction of ambiguously aligned reads at the first ~200 conserved positions for some *ospC* alleles (Supplemental Material S4).

Sensitivity of allele quantification is tested by generating a known proportion of simulated reads from two reference sequences. For example, a 10:1 mixed sample of short reads generated based on sequences of alleles “J” and “C” is quantified using the bioinformatics protocol, resulting in a ~13:1 quantification (Figure 2A).

**Figure 2.**
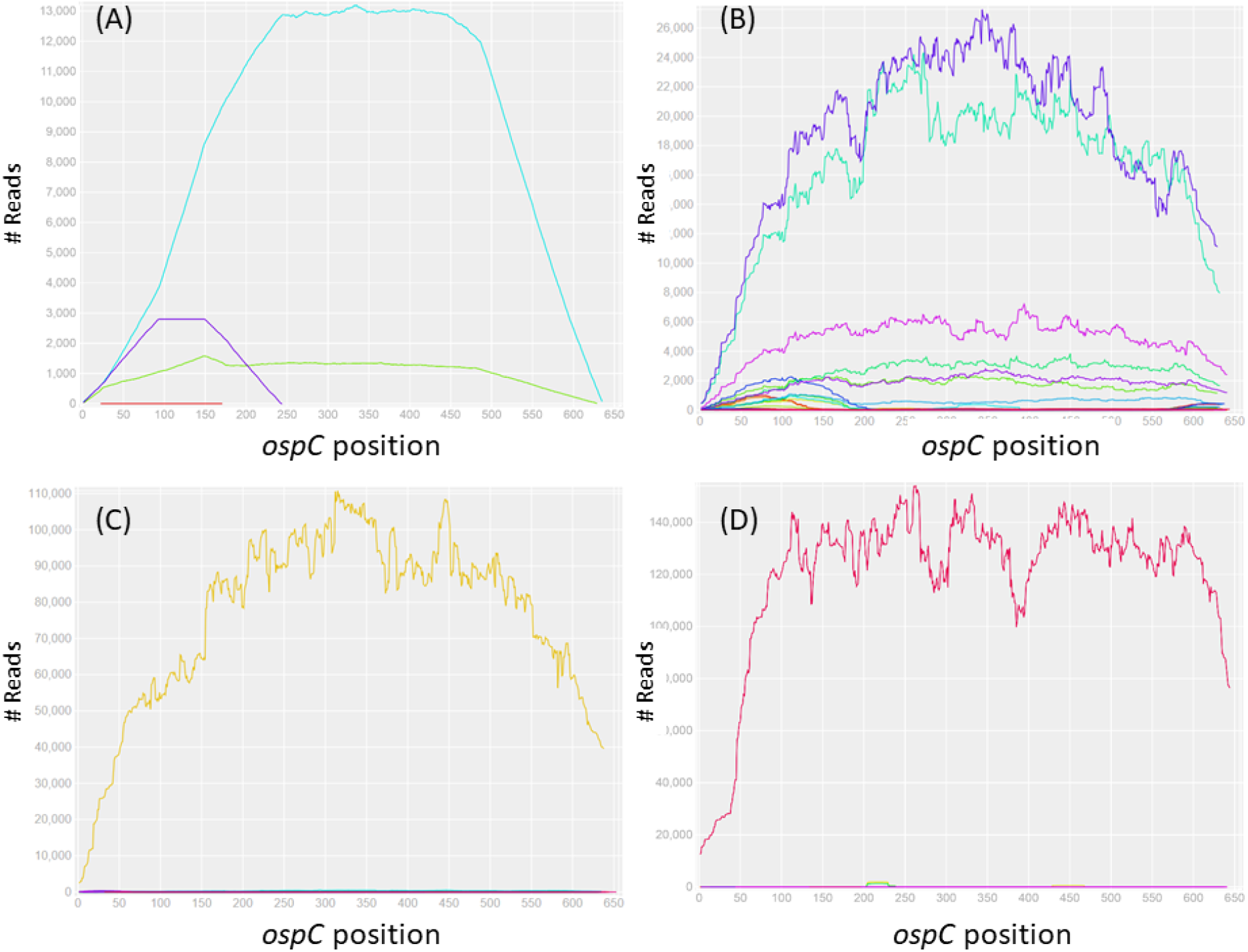
Read depths of *ospC* alleles in simulated (A) and tick (B, C, D) samples. (A) Simulated reads are generated based on nucleotide sequences of two *ospC* alleles (J and C) which are subsequently mixed in a 10:1 proportion. The reads are aligned to all 20 reference sequences (Supplemental Material S2) and only the two input alleles show complete, full-length coverages and approximately the same input proportion, validating the specificity and sensitivity of the bioinformatics protocol for allele identification (a full test of specificity is shown in Supplemental Material S4). (B) Seven *ospC* alleles (O, I, U, H, T, E, and K) are detected in an adult tick (#M119, male, RF, Fall 2016). (C) The universal *ospC* primer set is able to amplify not only the *ospC* locus in *Borreliella* species but also the *vsp* locus in *Borrelia miyamotoii* (see alignment in Supporting Material S1). Here the *vsp* locus is detected in a nymph tick (#N030, RF, Summer 2015). (D) A previously unknown *ospC* allele (“C14”, GenBank accession MH071431) is detected in a nymph tick (N150, RF, Summer 2016), suggesting presence of a new *B. bissettiae-* like species.

### New strain, spirochete load, and multiplicity

The NGS protocol is able to not only detect the presence of multiple strains but also quantify their relative frequency in individual ticks infected by multiple strains (Figure 2B). One allele (labeled as “C14_N150”) does not have known high-identity homologs in GenBank, with the top BLASTp hit as the *B. bissettiae* strain 25015 *ospC* with 75% identity in protein sequence (ACC45540) (Tilly et al., 1997). This allele likely represents an un-identified *Borreliella* species (Figure 2C). This allele was cloned, sequenced with Sanger method, and assigned a GenBank accession (MH071431). The full-length “F” allele was similarly cloned, sequenced with Sanger method, and assigned a GenBank accession (MH071432). The full-length “O” allele (MH071435) was sequenced with Sanger method directly from the singly-infected tick #N045 without cloning of the PCR amplicon. Sequences of full-length alleles “B3”, “N”, and “T” (MH071430, MH071433, and MH071436) were obtained by *de novo* assembly of short reads using metaSPAdes (Nurk et al., 2017). Our protocol is able to detect infection by *Borrelia miyamotoii*, as shown by the presence of one of its *vsp* (variable surface protein gene, locus name AXH25_04790) amplicons in samples (Figure 2D). The *vsp* allele was cloned, sequenced with Sanger method, and assigned a GenBank accession (MH071435).

Assuming that the spirochete load is correlated with total weight of PCR amplicons, we found that female adult ticks carry a significantly higher spirochete load than male adult ticks (*p*=0.022 by *t*-test), which in turn carry a higher infection load than nymphal ticks (*p*=8.1e-3) (Figure 3A). There is no significant difference in the average number of strains infecting a single tick (*p*>0.5 by Mann-Whitney test), although the median values are two strains per infected adult tick and one strain per nymphal tick (Figure 3B). Similarly, there are no significant differences in strain diversity measured by the Shannon diversity index between male, female, and nymphal ticks (*p*>0.5 by Mann-Whitney test; Figure 3C).

**Figure 3.**
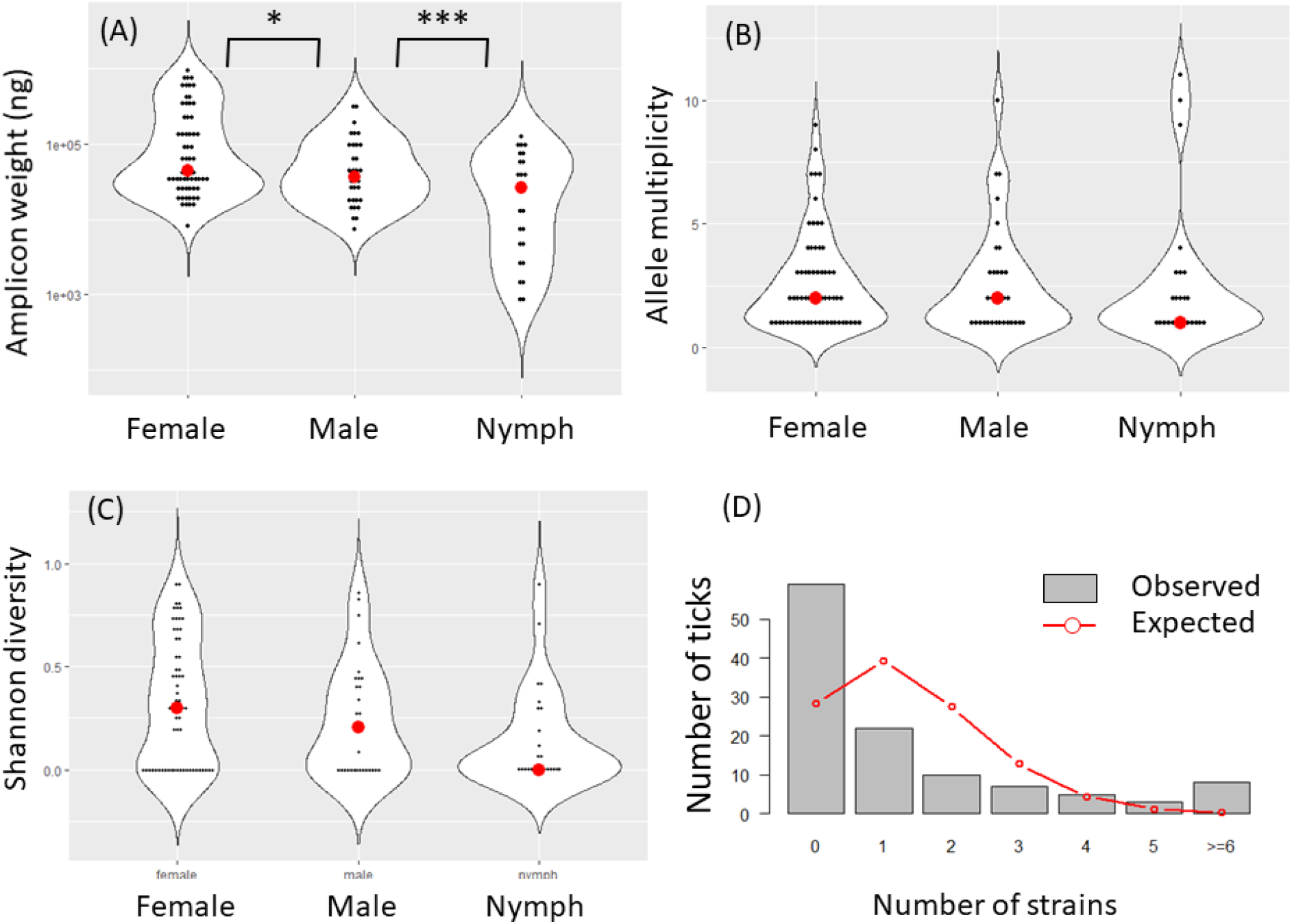
Spirochete load & diversity. (A) Spirochete loads, estimated with the weight of amplicons (*y*-axis, log10 scale), are significantly higher in female ticks than in males ticks, which in turn is higher than in nymphal ticks. (B) There is no significance differences among the three life stages in strain diversity as measured by the number of distinct strains within a tick (“multiplicity”). (C) Shannon diversity, which takes allele frequencies into account (see Material & Methods), is also not significantly different among the tick stages. These results support the notion that strain diversity in individual ticks is contributed more by a mixed inoculum in hosts than by the number of blood meals (Walter et al., 2016). (D) Observed counts of infected and uninfected adult ticks (N=144 from Sample #9, Figure 1). Expected counts are based on a Poisson model assuming that strains infect ticks independently. The observed distribution shows an over-abundance of uninfected ticks and ticks infected by five or more strains, while an under-abundance of ticks infected by 1-3 strains, suggesting reservoir hosts tend to be either uninfected or repeatedly infected.

### Aggregated infection & negative strain interactions

A previous study of multi-strain infection by *B. afzelii* in Europe found that strains tend to be aggregated in infected ticks, suggesting that infection of ticks and hosts is more successfully established by multiple spirochete strains than by a single strain alone (Andersson et al., 2013). Our data support their conclusion. In Sample #9, for example, we detected a total of 159 *ospC* alleles in 55 infected ticks out of a total of 114 processed adult ticks. Assuming a Poisson model of independent infection of individual strains with an average successful infection rate λ=159/114=1.395 strains per tick, we expect on average 28.2 uninfected ticks and 39.4 ticks infected by a single strain (the observed and expected counts are plotted in Figure 3D). In fact, 59 ticks are uninfected in this sample, more than twice the expected count. Meanwhile, 22 ticks are infected by a single strain, approximately half of the expected number. It appears that ticks tend to be either free of infection or infected by multiple spirochete strains, supporting the aggregated infection hypothesis (Andersson et al., 2013).

In infected ticks, previous studies conclude either a negative or a lack of interactions among co-infecting strains (Andersson et al., 2013; Durand et al., 2017; Walter et al., 2016). Our analysis supports presence of negative or inhibitory interactions among co-infecting strains. First, multiple strains tend to be unevenly distributed in their spirochete loads with some strains dominating others (e.g., Figure 2B). This is more generally shown with the Shannon diversity index, which is on average approximately half of the maximum attainable diversity (when all strains are equally abundant) in ticks with mixed infections (Supplemental Figure S5). There is, however, no evidence that any particular strains are consistently more dominant than other (Supplemental Figure S5). Second, when strains are independent from each other or facilitating each other’s growth, one expects ticks infected with multiple strains to have a higher spirochete load than ticks infected with a single strain. Conversely, if strains inhibit each other within a host or vector, one expects the total spirochete load to be either lower in ticks infected by multiple than by single strains or at similar levels. We plot the total pathogen load with respect to multiplicity or the Shannon diversity index in individual ticks (Figure 4). For the most part, the regression lines are not significantly different from a slope of zero except that the spirochete load in nymphs decreases significantly with increasing number of strains. The overall flat trend supports negative rather than facilitating interactions or a lack of any interactions between co-infecting strains (Durand et al., 2017; Walter et al., 2016).

**Figure 4.**
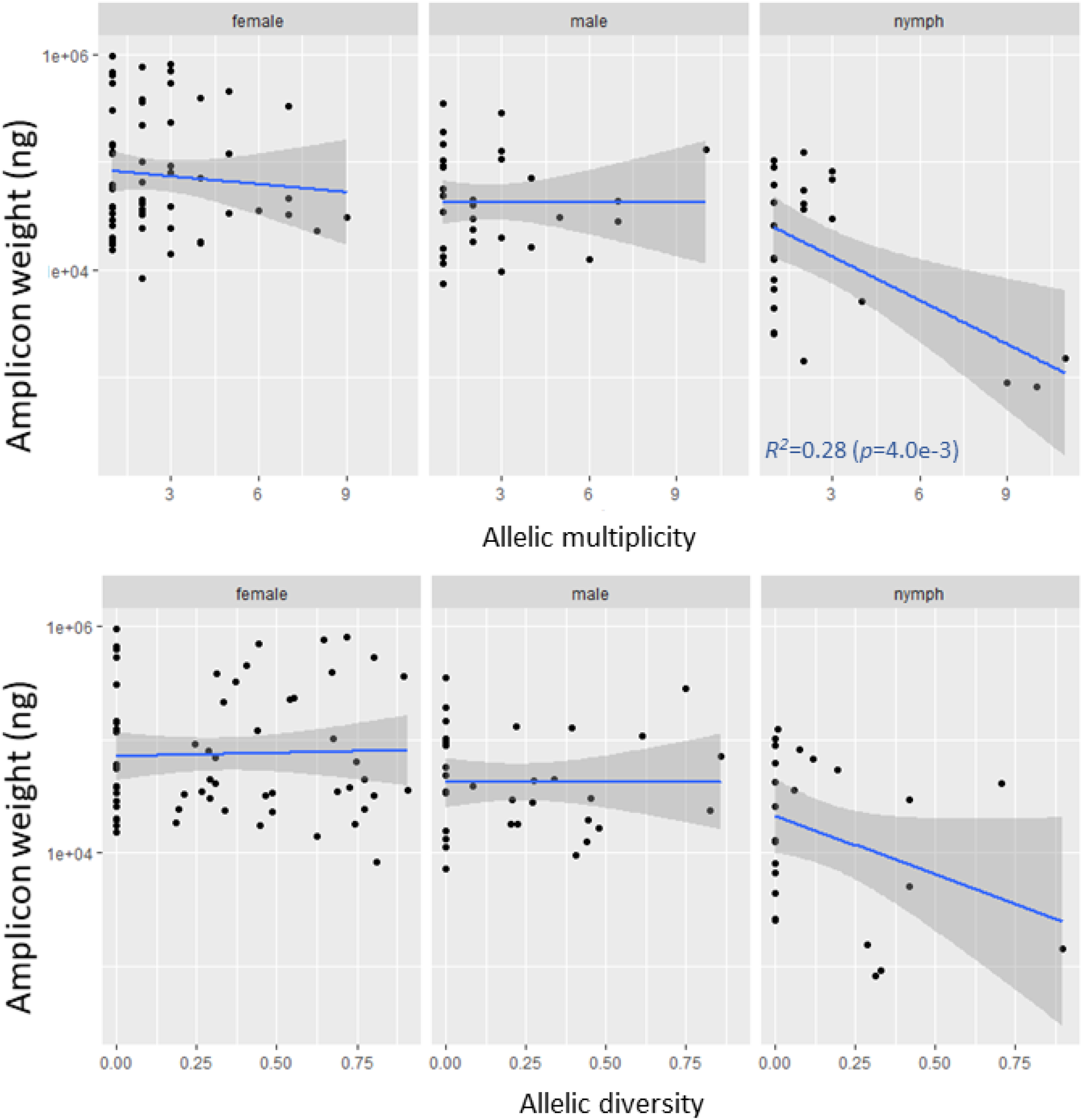
Negative interactions among co-infecting strains. (A) Spirochete load is either flat or decreasing with increasing strain multiplicity, supporting inhibitory interactions among co-infecting strains (Durand et al., 2017; Walter et al., 2016). (B) The pattern of inhibitory interactions holds when strain diversity is measured by Shannon diversity.

### Similar strain distributions among regions and life stages

Spirochete populations infecting adult and nymph ticks are similar in strain composition (*F_ST_*=4.7e-3 and *p*=0.089 by resampling, *p*=0.369 by *F*-test) (Figures 5A & 5C). Genetic differentiation between the Upper State and Long Island populations is more pronounced but nonetheless lacks statistical significance (*F_ST_*=5.3e-3 and *p*=0.052 by resampling, *p*=0.245 by *F*-test). The groups F and J strains appear to be more common on Long Island than Upper State while the group L strain shows the opposite pattern of distribution (Figures 5B & D).

**Figure 5.**
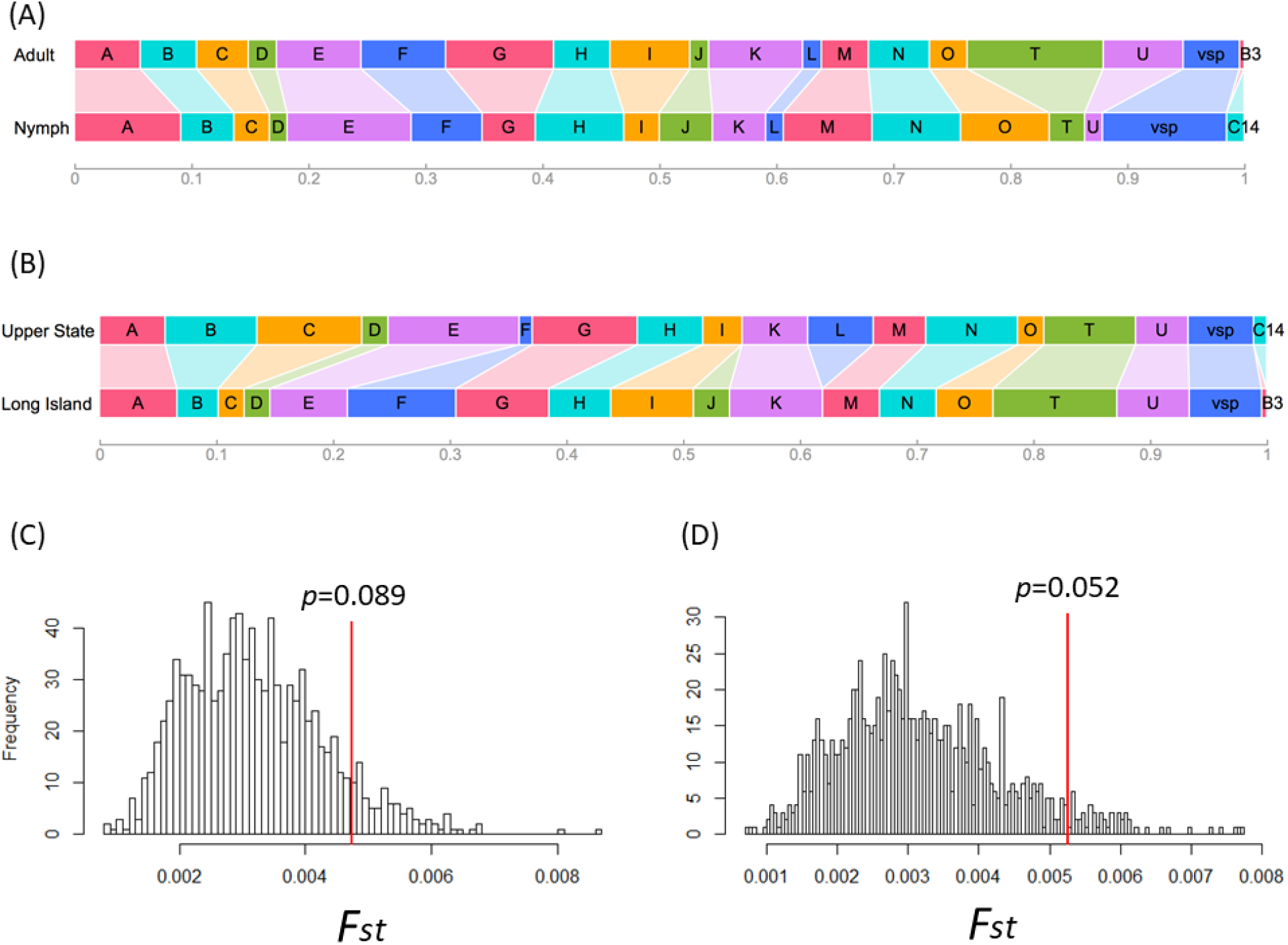
Geographic and life-stage differences in pathogen strain composition. (A) Strain composition (width of each colored rectangle representing frequency of an allele in a population sample) between those infecting adult ticks (Pop1+Pop3, see Table 1) and those infecting nymph ticks (Pop2 + Pop4). (B) Strain compositions in two regional populations (Upper State, Pop1 + Pop2; Long Island, Pop3 + Pop4;). (C) There is no significant genetic difference in strain composition between those infecting adult and those infecting nymph ticks (*p* value obtained by resampling 999 times; see Material & Methods). (D) Two regional populations show nonsignificant albeit stronger genetic differentiation.

## Discussion

In this report, we describe a new experimental and bioinformatics protocol for detecting and quantifying Lyme disease pathogen strains infecting individual ticks based on next-generation sequencing technology. Improving upon the previous Reverse-Line Blotting technology, the protocol allows *de novo* detection of previously unknown pathogen strains. Indeed, one of the ticks carries a putative new *Borreliella* species with a novel *ospC* allele (“C14”, MH071431). The protocol is highly sensitive and specific, enabling quantification of genetic diversity within single ticks and rigorously testing ecological hypotheses such as strain interactions (Figures 4) and genetic differentiation (Figure 5).

High-throughput sequencing has previously been used for quantification of Lyme strains in ticks based on either genome capture or *ospC* amplicons (Durand et al., 2017; Walter et al., 2016). Our technology is novel for an improved PCR protocol by using a set of universal PCR primers able to amplify full-length *ospC* in all *Borreliella* species as well as the *vsp* locus in *Borrelia miyamotoi*. Further, the PCR protocol is simplified from two rounds to a single round of thermal cycling. Due to these critical improvements in PCR protocol, our method could be readily used for detection and quantification of a broad range of Lyme disease (and possibly relapsing fever) pathogens in clinical and wildlife specimens across their species ranges worldwide. In addition, our technology is novel in using the Illumina short-read sequencing platform, in its supporting bioinformatics pipelines, and in its application to *Ixodes* ticks in North America.

Our strain identification method is based on the assumption of a strict one-to-one correspondence (i,e., complete genetic linkage) between *ospC* alleles and *B. burgdorferi* strains. The *ospC* locus is the most polymorphic single-copy locus in the *Borreliella* genome (Mongodin et al., 2013). The linkage between the *ospC* locus and the whole genome is indeed nearly complete for *Borreliella* populations in Northeast United States (Casjens et al., 2017; Mongodin et al., 2013). In fact, diversification of strains in local *Borreliella* populations is likely driven by frequency-dependent selection targeting the *ospC* locus (Haven et al., 2011; Qiu and Martin, 2014). However, linkage between *ospC* and other genomic loci is weaker in Midwestern and Southern US populations due to recombination and plasmid exchange (Hanincova et al., 2013; Mechai et al., 2015). Cross-species and cross-strain exchange of *ospC* alleles is also common in European populations. For example, whole genome sequencing showed that the European *B. burgdorferi* strain BOL26 obtained its *ospC* and its flanking genes from a con-specific strain through horizontal gene transfer (Qiu and Martin, 2014). For population samples elsewhere, therefore, it might be necessary to add a 2^nd^ locus for strain identification (Barbour and Cook, 2018). One complemental genetic marker could be the rRNA spacer *(rrs-rrlA)*, which is a single-copy and highly variable locus (Wang et al., 2014). Experimental methods for high-throughput sequencing of the *rrs-rrlA* locus however are yet to be developed.

While we are able to estimate relative spirochete loads in individual ticks based on quantification of *ospC* amplicons (Figure 2A), we have not attempted to directly quantify the number of spirochetes in infected ticks using methods such as quantitative PCR (Durand et al., 2017). In the future, we plan to quantify spirochete loads in individual ticks by running our experimental and bioinformatics procedures with known quantities of genomic DNA and generating a standard calibration curve.

Nonetheless, using relative estimates of spirochete loads in individual ticks, we are able to validate a number of hypotheses on multi-strain infections. First, the lack of differences in strain diversity between the questing adult ticks, which have taken two blood meals, and the questing nymphal ticks, which have taken one blood meal (Figure 3), supports the conclusion that strain diversity in individual ticks is for the most part due to mixed inoculum in infected hosts (Walter et al., 2016). Second, ticks are more likely to be infected by five or more strains than expected by chance (Figure 4D), supporting the aggregated infection hypothesis (Andersson et al., 2013). Rather than strains actively facilitating each other in establishing infections, however, strain aggregation in ticks may be a reflection of reservoir hosts being either free of spirochetes (in the case of resistant and healthy hosts) or infected by multiple strains (in the case of susceptible and weakened hosts). Regardless, it appears that once a host is infected by a strain, it becomes susceptible for super-infection by additional, immunologically distinct strains (Bhatia et al., 2018). Third, we found an uneven distribution of strains in infected ticks as well as a flat or decreasing spirochete load with increasing strain diversity (Figure 5), supporting inhibitory interactions among co-infecting strains driven by competitive growth in reservoir hosts (Durand et al., 2017; Walter et al., 2016). Fourth, we found weak genetic differentiation between populations from the two New York City suburbs (Figure 5B), suggesting either a recent common origin, or similar reservoir hosts, or both. Fifth, we observed co-circulation of *B. miyamotoii* and other *Borreliella* species in the same area. The lower prevalence as well as lower genetic diversity at *ospC* or *vsp* loci of these low-prevalence spirochetes relative to those of *B. burgdorferi* suggests that *ospC* hypervariability may be a key adaptation underlining the ecological success of *B. burgdorferi* in this region.

To summarize, we have established a next-generation sequencing-based, taxonomically broad procedure that has the potential to become a standard protocol for detecting and quantifying Lyme disease pathogens across the globe. The increased sensitivity of high-throughput sequencing technologies employed here and elsewhere highlights the prevalence of multiple infections in wildlife samples and a pressing need for broad spectrum vaccines for control and prevention of Lyme disease (Earnhart et al., 2007; Livey et al., 2011).

## Acknowledgements

We thank Ms Li Zhai and Dr Edward Skolnik of New York University School of Medicine for facilitating cloning experiments and Mr Brian Sulkow for participation of fieldwork. This work was supported by Public Health Service grants AI107955 (to WGQ) from the National Institute of Allergy and Infectious Diseases (NIAID) and the grant MD007599 (to Hunter College) from the National Institute on Minority Health and Health Disparities (NIMHD) of the National Institutes of Health (NIH) of the United States of America. The content of this manuscript is solely the responsibility of the authors and do not necessarily represent the official views of NIAID, NIMHD, or NIH.

## Supplemental Material

Text S1. Design of universal *ospC* primers for full-length amplification

Text S2. Nucleotide sequences of 20 *ospC* alleles used as references for strain identification

Text S3. Bioinformatics protocols

Figure S4. Specificity of allele identification

Figure S5. Strain distribution within infected ticks

